# The mechanism of homologous chromosome recognition and pairing facilitated by chromosome-tethered protein-RNA condensates

**DOI:** 10.1101/2023.12.24.573283

**Authors:** Da-Qiao Ding, Kasumi Okamasa, Yuriko Yoshimura, Atsushi Matsuda, Takaharu G. Yamamoto, Yasushi Hiraoka, Jun-ichi Nakayama

**Affiliations:** Advanced ICT Research Institute Kobe, National Institute of Information and Communications Technology, Kobe 651-2492, Japan; Division of Chromatin Regulation, National Institute for Basic Biology, Okazaki 444-8585, Japan; Graduate School of Frontier Biosciences, Osaka University, Suita 565-0871, Japan; Basic Biology Program, Graduate Institute for Advanced Studies, SOKENDAI, Okazaki 444-8585, Japan

**Author notes:** **Correspondence** Da-Qiao Ding, Advanced ICT Research Institute Kobe, National Institute of Information and Communications Technology, Kobe 651-2492, Japan., Jun-ichi Nakayama, Division of Chromatin Regulation, National Institute for Basic Biology, Okazaki 444-8585, Japan. **Funding information** MEXT | Japan Society for the Promotion of Science (JSPS), Grant/Award Number: JP19K06503, JP20H00454, JP23K05636, JP23H02462, JP20H03189, JP18H05532, JP19H03202 and JP20H05891 JST CREST Grant Number: JPMJCR2103 and JPMJCR22E2.

**Keywords:** Meiosis, homologous chromosome pairing, liquid-liquid phase separation, long noncoding RNA, *sme2* RNA associating protein, *in vitro* liquid-liquid phase separation assay

## Abstract

Pairing of homologous chromosomes during meiosis is crucial for successful sexual reproduction. Previous studies have shown that the fission yeast *sme2* RNA, a meiosis-specific long noncoding RNA (lncRNA), accumulates at the *sme2* locus and plays a key role in mediating robust pairing during meiosis. Several RNA-binding proteins accumulate at the *sme2* and other lncRNA gene loci in conjunction with the lncRNAs transcribed from these loci. These lncRNA-protein complexes form condensates that exhibit phase separation properties on chromosomes and are necessary for robust pairing of homologous chromosomes. To further understand the mechanisms by which phase separation affects homologous chromosome pairing, we conducted an *in vitro* phase separation assay with the *sme2* RNA-associated proteins (Smps) and RNAs. Our research has revealed that one of the Smps, Seb1, exhibits phase separation, which is enhanced by the addition of another Smp, Rhn1, and significantly increased by the addition of purified RNAs. Additionally, we have found that RNAs protect Smp condensates from treatment with 1,6-hexanediol. The Smp condensates containing different types of RNA display distinct FRAP profiles, and the Smp condensates containing the same type of RNA tend to fuse together more efficiently than those containing different types of RNA. Taken together, these results indicate that the RNA species in condensates determine their physical properties and suggest that regional RNA-Smp condensates with distinct properties ensure the pairing of homologous chromosomes.

## 1 INTRODUCTION

Sexual reproduction is achieved through a specific type of cell division called meiosis. This process produces gamete cells by reducing the diploid set of homologous chromosomes in the progenitor cells to a haploid set, through two rounds of consecutive chromosome segregation. One of the key processes during meiosis is the recombination of homologous chromosomes that come from each parent cell. Recognition and pairing of homologous chromosomes are essential prerequisites for this process of homologous recombination to take place. In contrast to the highly conserved process of recombination, the process of homologous chromosome pairing varies greatly among different organisms, and there is currently no unified theory to explain the underlying mechanism of homologous chromosome recognition. One relatively conserved process of pairing is the formation of a chromosome bouquet, in which telomeres of all chromosomes cluster in a relatively small area of the nuclear envelope.^1,2^ This bouquet configuration effectively aligns homologous chromosomes in the same direction and reduces the spatial distances between them.^3,4^

Previously, we found that in the fission yeast *Schizosaccharomyces pombe, sme2* RNA, a meiosis-specific long noncoding RNA (lncRNA), accumulates at the *sme2* gene locus in meiotic prophase and mediates a recombination-independent robust pairing.^5^ We further identified several conserved RNA-binding proteins that are associated with the *sme2* RNA (*sme2* RNA-associated proteins: Smps); the Smps are required for the lncRNA dot integrity and robust pairing of homologous chromosomes.^6^ These proteins accumulate as distinct dots on chromosomes, at the *sme2* and two other chromosomal loci, together with meiosis-specific lncRNAs transcribed from these loci. The lncRNA–Smp complexes exhibit liquid-liquid phase separation (LLPS) properties, since 1,6-hexanediol treatment reversibly disassembled these lncRNA-protein condensates and disrupted the pairing of the associated loci. smFISH (single molecular RNA fluorescence in situ hybridization) analysis revealed that droplet-like RNA-protein condensates on each homologous locus fuse together to form a single focus connecting the juxtaposed homologous chromosomes.^6^ When homologous loci were manipulated to express different types of RNA, these RNA droplets could not fuse with each other,^6^ further implying that RNA plays a key role in distinguishing the homology during chromosome recognition and pairing.

LLPS has been widely recognized as the principal mechanism for the formation of membrane-less organelles and biomolecular condensates in cells.^7^ Biochemical interactions and processes of LLPS associated with proteins, RNA, and DNA have been increasingly identified. Many nuclear condensates are associated with specific chromosomal loci, for example, nucleolar organizing region, transcription factories at enhancers, HP1α foci at heterochromatin, and Shelterin at telomeric chromatin.^8-11^ The Smp-lncRNA condensates with LLPS properties may play a novel role in joining two homologous chromosomes during meiotic prophase. To explore the mechanism of Smp-lncRNA condensate formation and its role in homologous chromosome pairing, we here performed *in vitro* LLPS assays for representative Smps and RNAs. We investigate whether the Smp components can form phase-separated liquid droplets *in vitro*, and how other components affect this process. Furthermore, we attempt to identify the key components that determine the specificity of homologous recognition during pairing.

## 2 MATERIALS AND METHODS

### 2.1 Preparation of recombinant proteins

The coding sequence for the maltose binding protein (MBP) with a recognition site for 3C protease was amplified by PCR and cloned into pFastBac-HTb (Invitrogen) (pFastBac-HTb-MBP-3C). The coding sequence for mEGFP or mCherry with a recognition site for TEV protease was amplified by PCR and cloned into the pFastBac-HTb-MBP-3C (pFastBac-HTb-MBP-3C-mEGFP/mCherry-TEV). The coding sequences for Seb1 and Rhn1 were then amplified by PCR using *S. pombe* genomic DNA or cDNA and subcloned downstream of the TEV site of the pFastBac-HTb-MBP-3C-mEGFP/mCherry-TEV vector. Recombinant MBP-mEGFP/mCherry-Seb1 and MBP-mEGFP/mCherry-Rhn1 were expressed in Sf9 cells using the Bac-to-Bac baculovirus expression system (Invitrogen), in accordance with the manufacturer’s instructions. Sf9 cells harvested from 250 ml culture were lysed in 40 ml of lysis buffer [50 mM Tris-HCl (pH7.5), 500 mM KCl, 5% glycerol, 5 mM imidazole] containing 1 mM phenylmethylsulfonyl fluoride (PMSF) and protease inhibitor cocktail (Complete™; Roche), and recombinant proteins were purified by immobilized metal affinity chromatography (TALON; Clonetech) in accordance with the manufacturer’s instructions. The eluted proteins were pooled, dialyzed against dialysis buffer [50 mM Tris-HCl (pH7.5), 1 M NaCl, 1 mM EDTA, 10% glycerol, 1 mM dithiothreitol (DTT)], frozen in LN_2_, and stored at −80°C before further purification. To remove the MBP moiety, purified MBP-mEGFP/mCherry-Seb1 and MBP-mEGFP/mCherry-Rhn1 proteins were incubated with HRV 3C protease (Takara; 1 U/1 g of target recombinant proteins) for 1 h at room temperature, and reaction samples were loaded onto 1 ml of amylose resin (NEB). Proteins in the flow-through fractions were further purified by gel filtration chromatography (Superdex 200pg; Cytiva), and proteins in the eluted fractions were pooled, concentrated using a centrifugal filter unit (Amicon Ultra-4; Merck), frozen in LN_2_, and stored at −80°C before use.

### 2.2 Preparation of RNA

RNAs of *sme2, omt3, gfp* and *rhn1* were prepared using an *in vitro* transcription T7 kit (Takara No.6140) and purified following the manufacturer’s instructions. Primers are listed in Table 1. Fluorescently labeled RNA was obtained by replacing 1/4 volume of CTP with Cyanine 3-CTP (Cy3-CTP) or Cyanine 5-CTP (Cy5-CTP) (PerkinElmer NEL581001EA) in the *in vitro* transcription reaction.

**Table 1.**
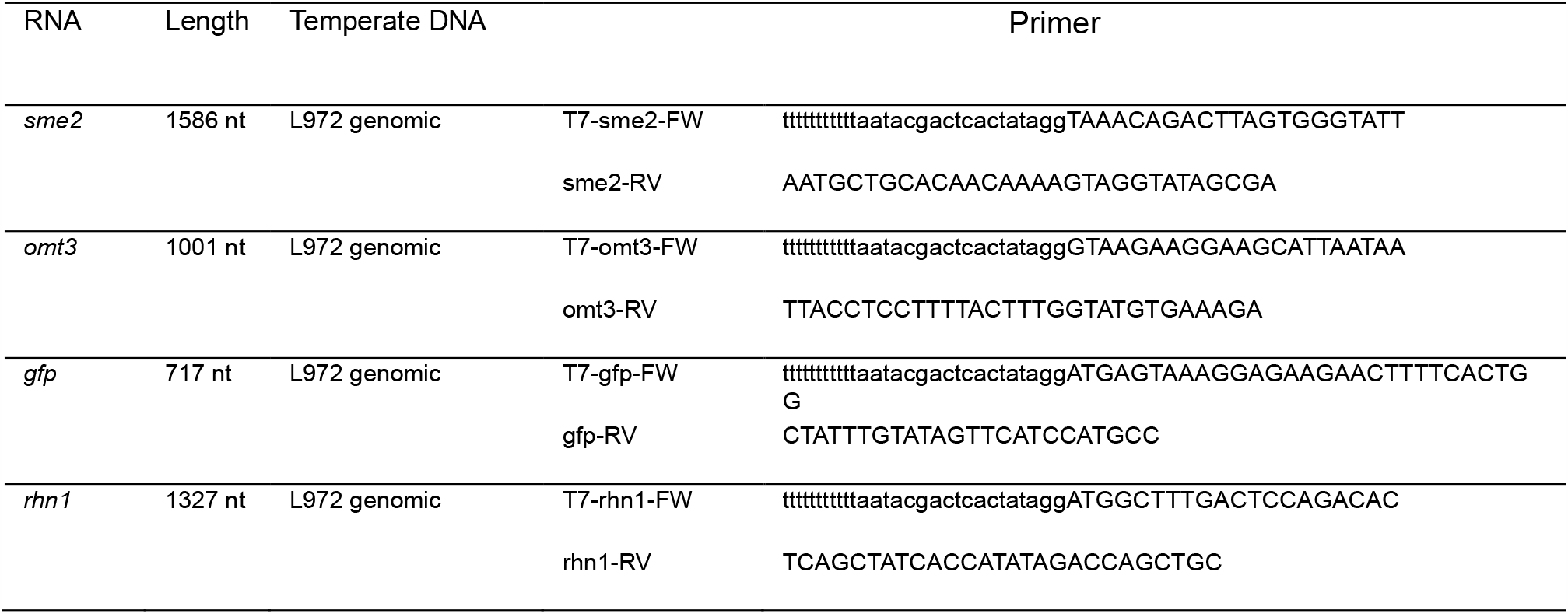
List of primers.

### 2.3 LLPS experiment

LLPS of Seb1 and other proteins was induced in the presence of 10% polyethylene glycol (PEG4000, Wako, 162-09115) as a crowding agent. In our experience, droplets formation of Seb1 protein never occurs without PEG or the alternative crowding agent Dextran (Nacalai Tesque 10911-44). The proteins were mixed with the phase separation buffer [20 mM HEPES-NaOH (pH7.0), 10% PEG4000, 150 mM NaCl] ^12^ and incubated for 30 min at room temperature (23 ± 2°C), and then placed on a lipid-coated imaging chamber for microscopic observation. The lipid-coated imaging chamber was set up following the steps outlined in a previously published protocol.^13^ A coverslip (18 × 18 mm) attached to three layers of double-sided tape (0.01 mm thickness, Teraoka Seisakujou No. 7070) was used to form an 18 × 9 × 0.03 mm rectangular chamber with two openings on the opposite sides on a large coverslip (24 × 60 mm) for observation.

### 2.4 Microscopic observation and FRAP experiment

For deconvolution microscopy, a DeltaVision Elite microscope (Cytiva) with an objective lens 60× PlanApo NA 1.4 Oil SC (Olympus) set up in a temperature-controlled room at 26°C was used. For quantitation of droplet formation, 3D image stacks for 20 to 30 areas (1024 × 1024 pixels, with the image pixel size of 0.1075 × 0.1075 μm) were taken. Data analysis was carried out by using SoftWoRx software on the DeltaVision system. The area and number of droplets were estimated using the 2D Polygon Finder function of the software.

Fluorescence recovery after photobleaching (FRAP) experiments were conducted using the Photokinetics function of a DeltaVision|OMX microscope SR (Cytiva) with an objective lens 60x PLAPON NA1.42 Oil (Olympus). We used a 488 nm laser, 10% transmission, and duration of 0.01 sec to bleach the GFP fluorescence signal in an entire droplet. In all of the FRAP experiments, those droplets with a diameter of 1.0 ± 0.2 μm were selected for photobleaching. FRAP data analysis was performed by using the PK analysis function of the softWoRx software with the two components model. In each experiment, FRAP data from 10 droplets were collected.

## 3 RESULTS

### 3.1 Smps contain low-complexity region in their sequences

We have identified six Smps that co-localize in foci at the *sme2* gene locus and are involved in *sme2* RNA-mediated homologous pairing ^6^ as listed in Figure 1A. The Smp foci disappear upon the addition of 1,6-hexanediol and reappear after its removal, suggesting that they are protein-RNA condensates formed through LLPS. ^6^ Since intrinsically disordered regions (IDRs) are characteristic of proteins for LLPS, we surveyed IDRs in Smps by using the PSIPRED Workbench established by The Bioinformatics Group at University College London (http://bioinf.cs.ucl.ac.uk/psipred/). As shown in Figure 1B, approximately half of Seb1 is predicted to be IDR with a confidence score higher than 0.5. Some other Smps also share a similar feature of high enrichment of IDRs (Figure 1A). Since Seb1 is the only protein that has an RNA binding domain (PomBase https://www.pombase.org)^14^ and physical interaction between Seb1 and Rhn1 has been detected,^15^ we focused our *in vitro* study on these two proteins.

**Figure 1.**
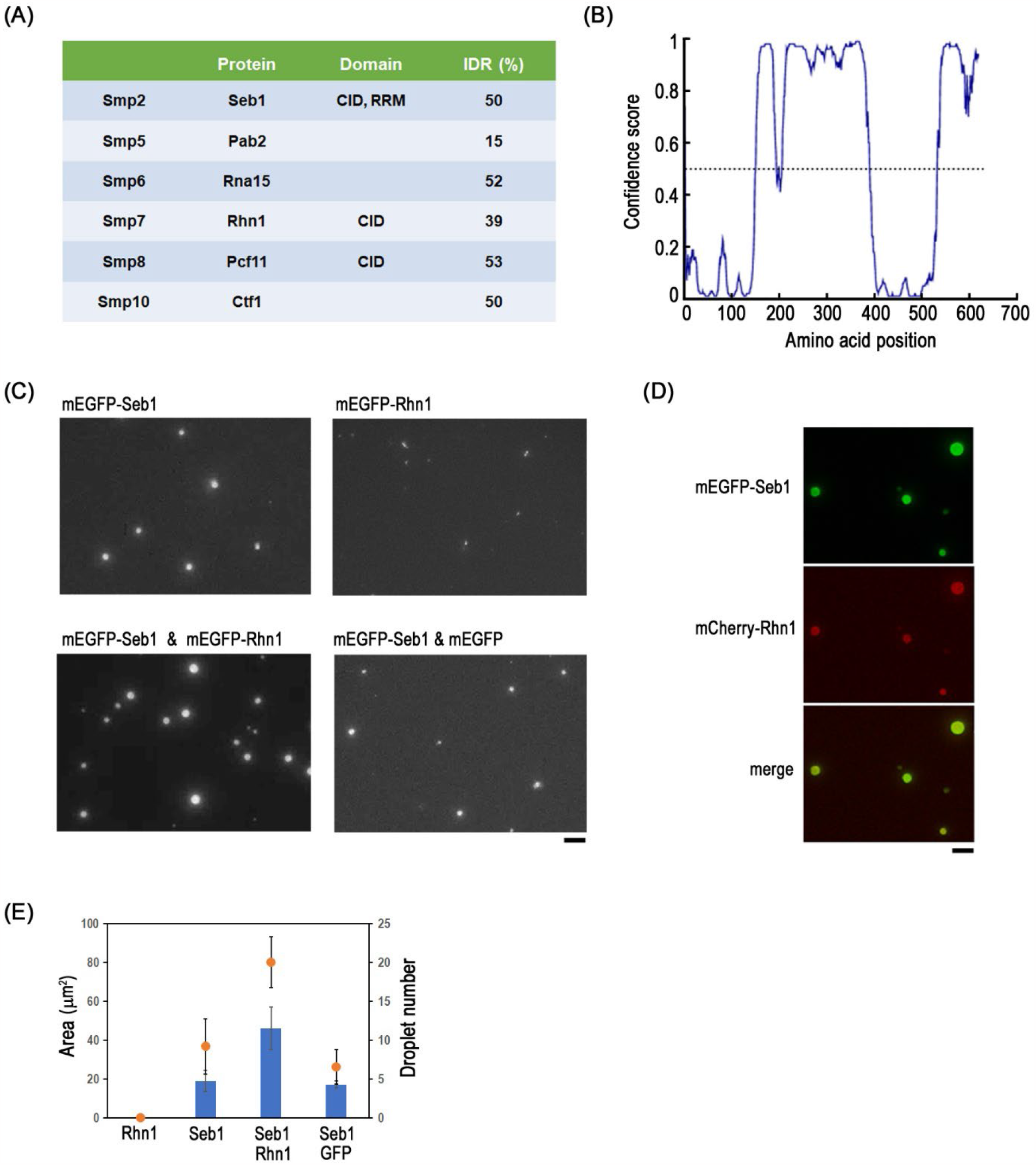
Seb1 protein forms phase-separated droplets *in vitro*. (A) Predicted domain and percentage of IDR region in Smps. (B) Predicted domain of IDR in Seb1. (C) Representative images of droplet formation of mEGFP-Seb1, mEGFP-Rhn1 (upper panels), mixture of mEGFP-Seb1 and mEGFP-Rhn1, and mixture of mEGFP-Seb1 and mEGFP (lower panels). Concentration for all proteins was 1 μM. (D) Representative image of mCherry-Rhn1 accumulated in mEGFP-Seb1 droplets. Bars, 5 μm. (E) Quantitation of the total droplet areas (blue bars) and droplet numbers (orange circles) in a 1024 × 1024 pixel microscopic image area (13,083 μm^2^). Means of three independent experiments are presented. Error bars show standard deviations.

### 3.2 Seb1 protein forms phase-separated droplets *in vitro*

We used a baculovirus expression system and insect cells to obtain recombinant proteins (Supplementary Figure 1, see also Materials and Methods). Purified Seb1 and Rhn1 proteins fused with mEGFP or mCherry were used for an *in vitro* LLPS assay (hereafter, we refer to these proteins simply as Seb1 and Rhn1, respectively, unless otherwise indicated). After the proteins were mixed in the phase separation buffer and incubated for 30 min, we observed phase-separated droplet formation using a fluorescence microscope. As shown in Figure 1C (upper left panel), when purified Seb1 protein (concentration of 1 μM) was mixed with a phase separation buffer, droplets ranging from 0.3 to 4.4 μm in diameter were observed. However, droplet formation was not observed when purified Rhn1 was used under the same experimental conditions (Figure 1C, upper right panel), only a few small aggregates (not round shape) were observed. In addition, no droplet formation was found when purified mEGFP or mCherry was mixed with a phase separation buffer (Supplementary Figure 2A). The number of droplets formed was counted; the quantitative data on droplet formation are presented in Figure 1E. When both Seb1 and Rhn1 were added to the phase separation buffer simultaneously, more droplets formed compared to when only Seb1 was used (Figure 1C, lower left panel; Figure 1E). Additionally, Rhn1 was found to be concentrated within the droplets formed by Seb1 (Figure 1D). In contrast, when mEGFP was added instead of mEGFP-Rhn1, no significant effect on droplet formation was observed (Figure 1C, lower right panel; Figure 1E). These results suggest that Seb1 is capable of undergoing LLPS and that Rhn1 promotes LLPS formation by Seb1.

### 3.3 *sme2* RNA promotes the phase separation of Seb1

As previously demonstrated, the formation of Smp dots *in vivo* requires the presence of RNA transcribed and tethered to chromosomes.^6^ To further investigate the effects of RNA on LLPS of Seb1, we added *in vitro*-transcribed and purified *sme2* lncRNA to the mixture of Seb1 (with or without Rhn1) in the phase separation buffer. As shown in Figure 2A and B, the addition of *sme2* lncRNA resulted in the formation of larger droplets of Seb1 than for the protein alone. The addition of *sme2* lncRNA also increased the number of droplets formed (Figure 2B). Notably, droplet formation was not observed when only *sme2* lncRNA was mixed with the phase separation buffer (Supplementary Figure 2B). Nevertheless, the *sme2* lncRNA was found to be concentrated within the Seb1 droplets (Figure 2C). The LLPS of Seb1 was effectively promoted with *sme2* lncRNA at a low concentration of 12.5 nM, increasing the concentration beyond that level did not result in a proportional increase in LLPS (Figure 2D). In conclusion, these results suggest that, while *sme2* lncRNA itself does not possess the ability to undergo LLPS, it effectively promotes the LLPS of Seb1 at the nanomolar concentration level.

**Figure 2.**
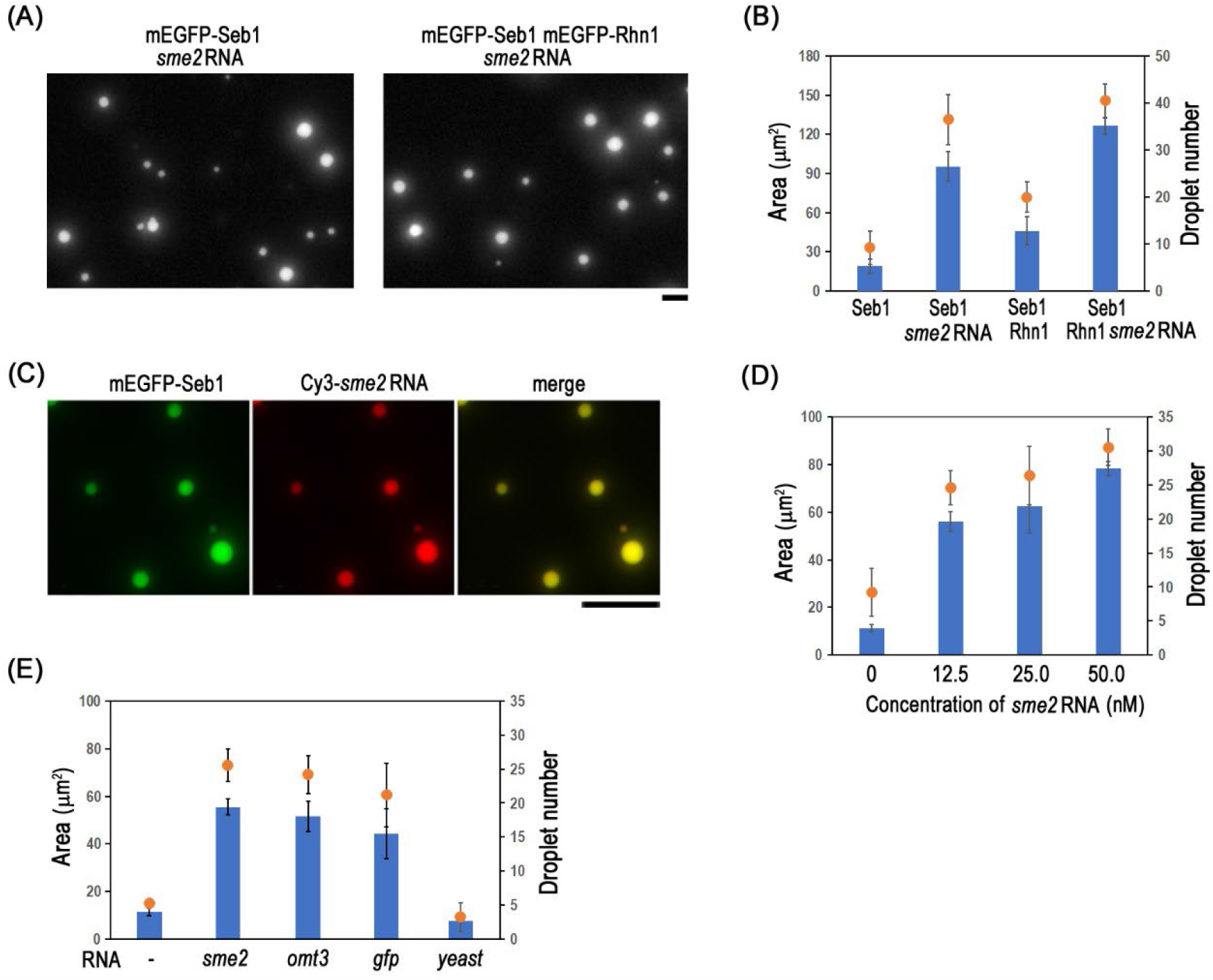
*Sme2* RNA promotes the phase separation of Seb1. (A) Representative images of droplet formation of mEGFP-Seb1 (without or with Rhn1) when 12.5 nM *sme2* RNA was added. Concentration for all proteins was 1 μM. (B) Quantitation of the total droplet areas (blue bars) and droplet numbers (orange circles) in various experimental conditions. Averages of three independent experiments are presented. Error bars show standard deviations. Droplet formation of 1 μM mEGFP-Seb1 (without or with 1 μM Rhn1) when 12.5 nM *sme2* RNA was added. (C) Representative image of Cy3-*sme2* RNA accumulated in mEGFP-Seb1 droplets. Bars, 5 μm. (D) and (E) Quantitation of the total droplet areas (blue bars) and droplet numbers (orange circles) in various experimental conditions. Means of three independent experiments are presented. Error bars show standard deviations. Droplet formation of mEGFP-Seb1 and mEGFP-Rhn1 (0.5 μM each) when *sme2* RNA concentration was increased (D). Comparison of droplet formation of mEGFP-Seb1 and mEGFP-Rhn1 (0.5 μM each) when various kinds of RNA (12.5 nM) were added (E).

To determine whether the effect of *sme2* lncRNA on Seb1 LLPS promotion is limited to *sme2* lncRNA, we tested other coding and noncoding RNAs. As shown in Figure 2E, both lncRNA (*omt3*) and protein-coding RNAs (*gfp, rhn1*) had similar LLPS-promoting effects as *sme2* lncRNA. However, when yeast total RNA, which mainly consists of ribosomal RNAs, was added, it did not have an effect similar to that of *sme2* lncRNA (Figure 2E). These results suggest that not only *sme2* lncRNA, but also other lncRNAs or protein-coding RNAs, can promote Seb1 LLPS. However, structural RNAs such as rRNA may not be effective in mediating LLPS by Smps such as Seb1 and Rhn1.

### 3.4 RNA enhances the resistance of Seb1 droplets to 1,6-hexanediol treatment

To investigate the role of RNA in promoting LLPS of Seb1, we conducted experiments by treating Seb1 droplets with 5% 1,6-hexanediol, in either the presence or the absence of RNA. As shown in Figure 3A (upper panel), in the absence of RNA the 1,6-hexanediol treatment of Seb1 droplets caused shrinkage, forming small pin dots. However, in the presence of *sme2* lncRNA, the droplets are largely maintained their size (Figure 3A, lower panels). The area of the droplets before and after the treatment was quantified, as shown in Figure 3B. The droplet area showed a 70% decrease without RNA, but only a 30% decrease with the addition of *sme2* lncRNA. The *gfp* RNA also had a stabilizing effect on the droplets, but to a lesser extent than the *sme2* lncRNA (Figure 3B). These results indicate that RNAs act as a scaffold within Seb1 droplets, reinforcing and stabilizing the droplet structures.

**Figure 3.**
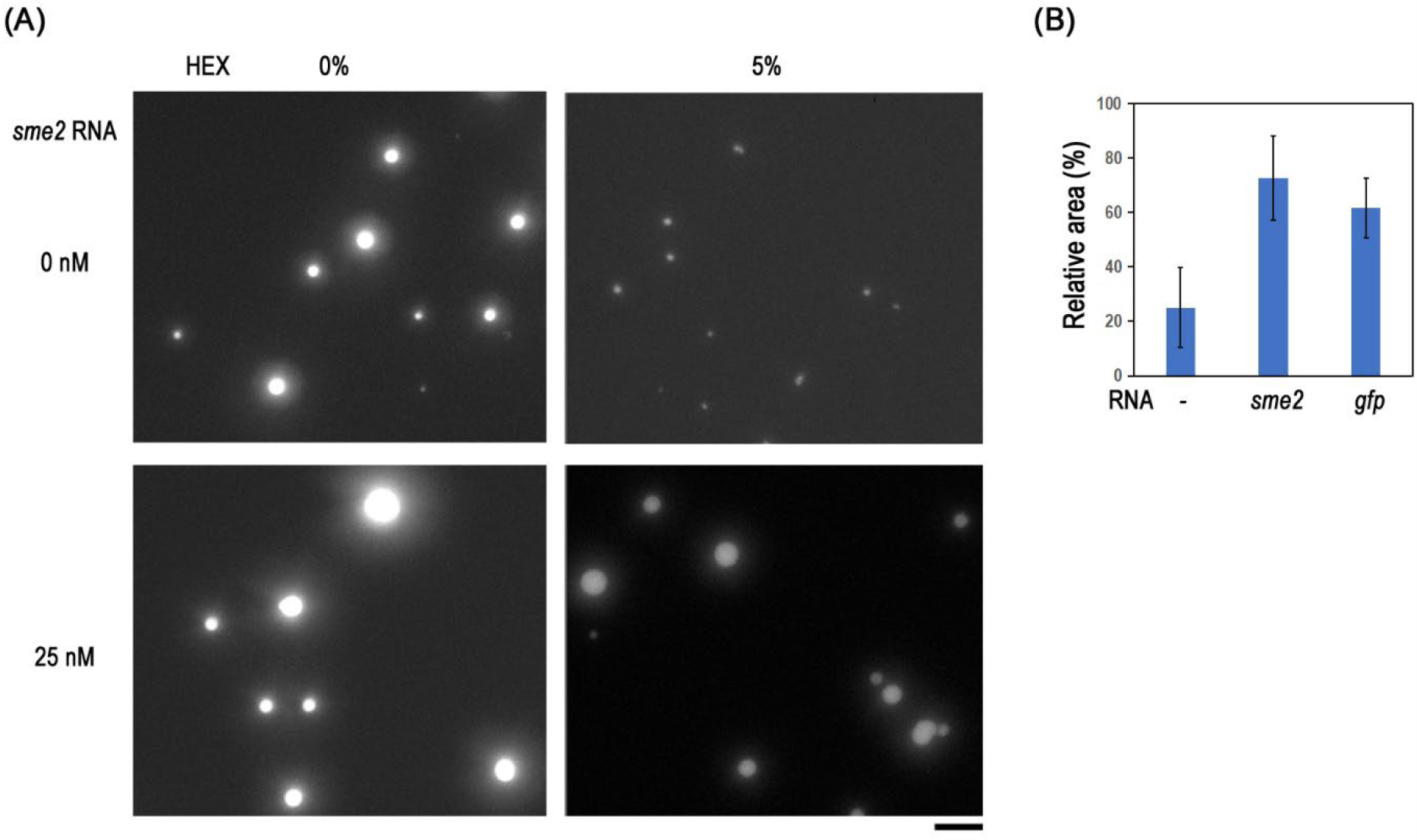
RNAs enhance the resistance of Seb1 droplets to 1,6-hexanediol treatment. (A) Representative images of mEGFP-Seb1/Rhn1 droplets after treatment with 5% 1,6-hexanediol (HEX) in the absence (upper panel) and presence (lower panel) of 25 nM *sme2* RNA. Bar, 5 μm. (B) Relative droplet areas (blue bars) compared to no 1,6-hexanediol treatment in the absence or presence of *sme2* or *gfp* RNA. Means of three independent experiments are presented; error bars show standard deviations.

### 3.5 RNA reduces the dynamic movement of molecules within liquid droplets

To further elucidate the role of RNA in enhancing Seb1 droplet formation and tolerance to 1,6-hexandiol treatment, we conducted FRAP experiments to compare the dynamics of protein exchange between the inside and outside of the droplets in the presence or absence of RNA. Representative examples of FRAP in mEGFP-Seb1 droplets are shown in Figure 4A. The recovery of fluorescence was slower with *sme2* lncRNA than that without RNA. The results of FRAP with different concentrations of *sme2* lncRNA are shown in Figure 4B. Slower recovery of fluorescence was found at *sme2* lncRNA concentrations of 6.3 and 12.5 nM, but not at 3.1 nM, and higher RNA concentrations resulted in slower recovery. We also performed FRAP experiments with different RNAs, namely, noncoding (*omt3*) and coding ones (*gfp, rhn1*) RNAs. All of the RNAs resulted in slower recovery than in the absence of RNA, and each RNA showed a specific FRAP profile (Figure 4C). These results suggest that the RNA species and concentrations can change the biophysical properties of the Seb1 droplets.

**Figure 4.**
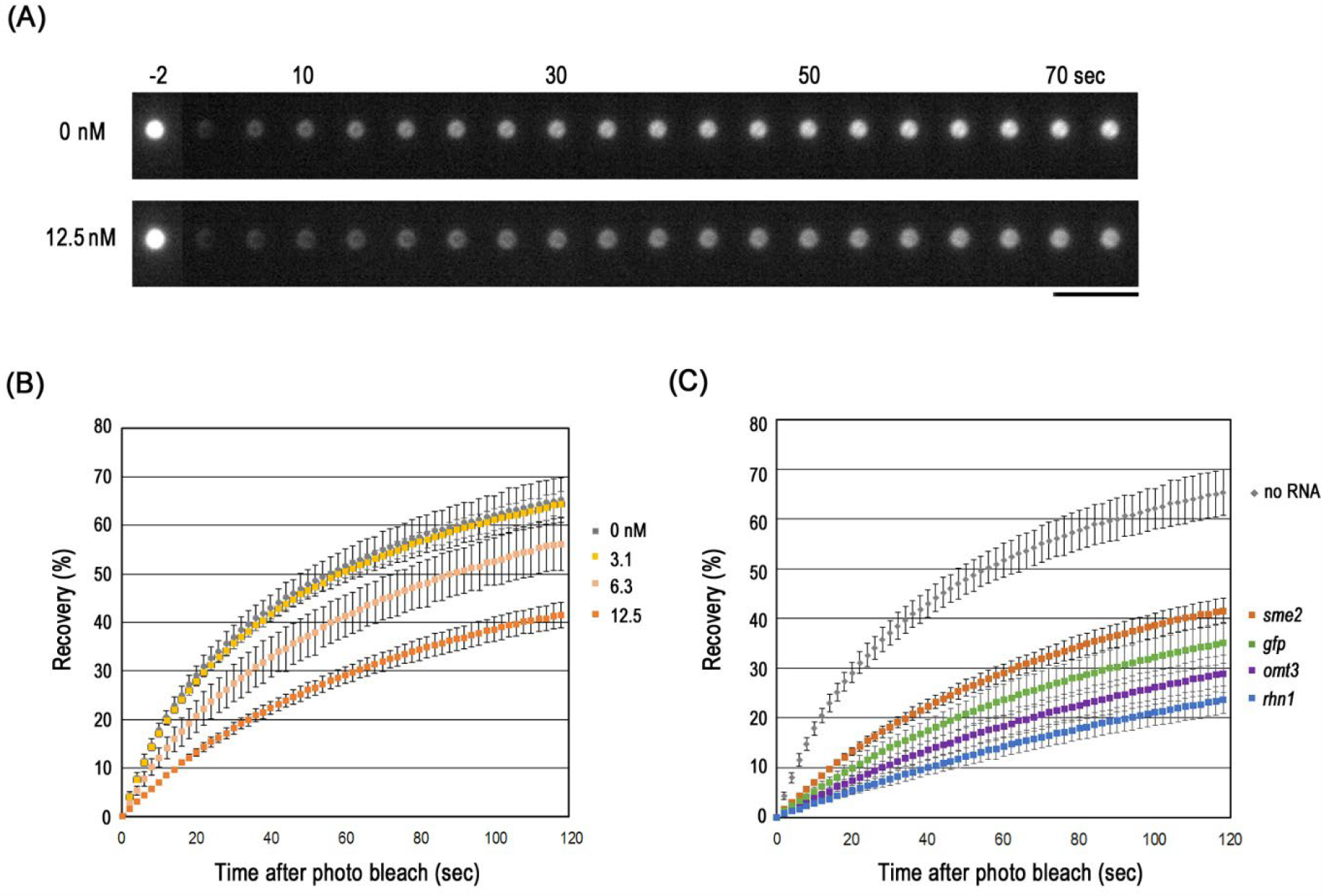
RNAs reduce the dynamic movement of molecules within liquid droplets. (A) Representative time lapse images of FRAP experiments of mEGFP-Seb1/Rhn1 droplets in the absence (upper panel) and presence (lower panel) of 12.5 nM *sme2* RNA. Bar, 5 μm. (B) (C) Recovery of mEGFP-Seb1 fluorescence with different concentrations of *sme2* RNA (B) or different kinds of RNA (all at a concentration of 12.5 nM) in the droplets (C). Error bars show standard deviations from three independent experiments.

### 3.6 RNA species in the Smp/RNA droplets modulate the efficiency of droplet-droplet fusion

We hypothesized that Smp/RNA droplets with similar biophysical properties could easily fuse with each other, thus providing the basis for the recognition and pairing of homologous chromosomes.^6^ To test this idea, we mixed two types of pre-prepared Seb1 droplets containing different RNAs; one of the RNAs was unlabeled and the other was labeled with Cy5-CTP (Figure 5A). In the mixture of droplets, we counted the number of droplets in which no Cy5 fluorescence could be detected (indicated by arrows in Figure 5A, B). If no droplet fusion occurs, about 50% of the droplets should have no Cy5 fluorescence staining. The decrease in the percentage of unlabeled RNA droplets is proportional to the fusion between labeled and unlabeled droplets. The results showed that the percentage of Seb1 droplets with unlabeled *sme2* lncRNA was lower in the experiments using droplets containing Cy5-*sme2* RNA than in those using droplets containing Cy5-*omt3* RNA (Figure 5C). This suggests that, under the present experimental conditions, Seb1 droplets containing *sme2* lncRNA have a higher tendency to fuse with droplets that also contain *sme2* lncRNA than those containing different RNA species such as *omt3* lncRNA.

**Figure 5.**
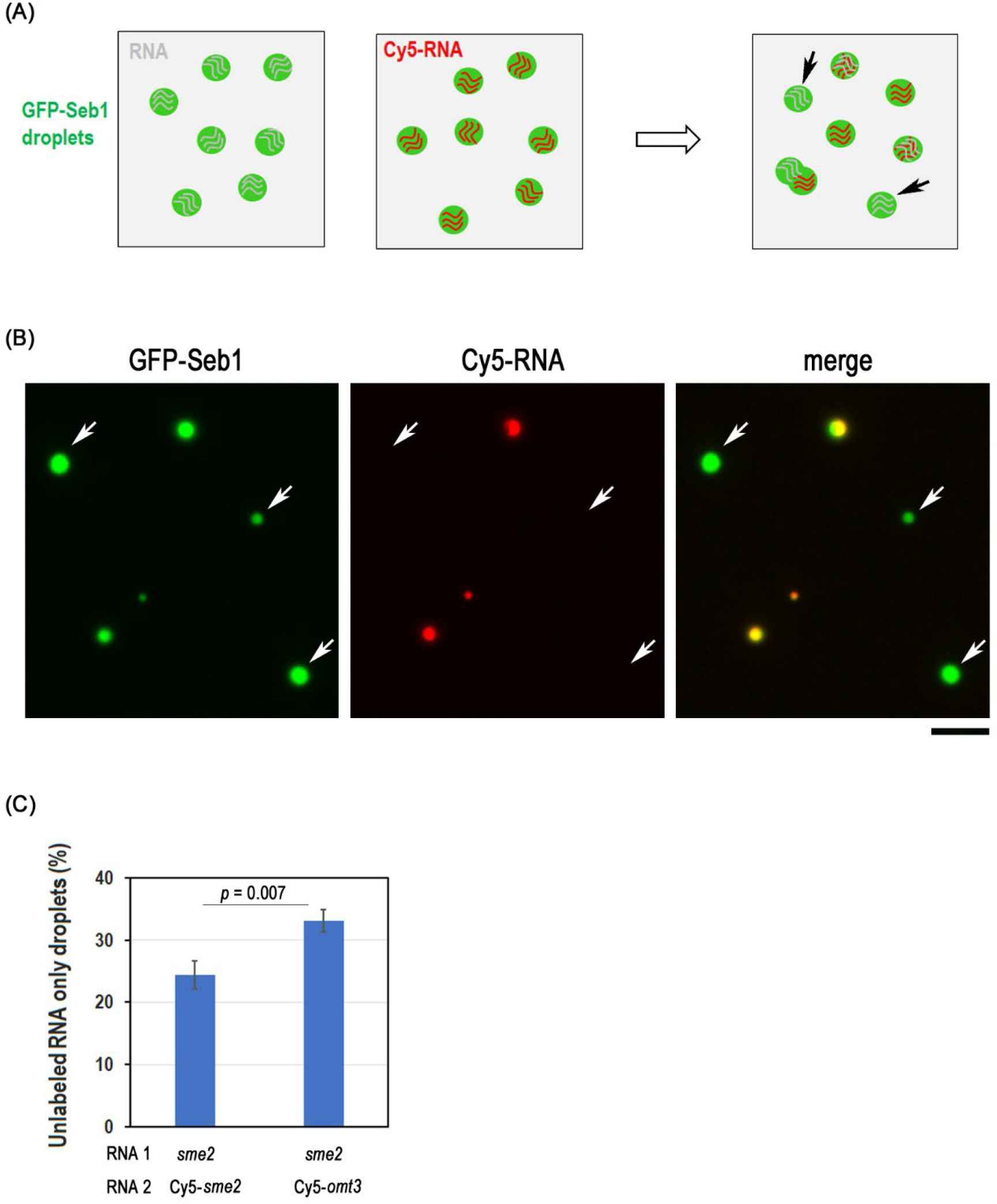
Smp droplets containing *sme2* RNA have a higher tendency to fuse with droplets that also contain *sme2* RNA. (A) Schematic of the design of the experiment. Equal volume of two types of pre-prepared Seb1 droplets containing different RNAs; one with unlabeled RNAs and the other with Cy5 labeled RNA, are mixed. When droplets containing different type of RNA fuse, number of unlabeled droplet will decrease. The number of unlabeled RNA containing droplets (arrows) in the mixture are counted. (B) Representative images of a mixture of mEGFP-Seb1 droplets (green) which containing unlabeled *sme2* RNA (as indicated by arrows) or Cy5 labeled *sme2* RNA (red). (C) Quantitation of the percentage of droplets containing only unlabeled *sme2* RNA. Error bars show standard deviations from three independent experiments. Student’s *t*-test was used for the statistical analysis.

## 4 DISCUSSION

In this study, we have demonstrated the ability of one of the Smps, Seb1, to undergo LLPS *in vitro*. Both noncoding and coding RNAs play a significant role in enhancing the LLPS of Seb1 by increasing the stability of the resulting droplets. Additionally, each type of RNA imparts a distinct biophysical property to these droplets. Notably, droplets containing the same RNA molecules exhibit a higher tendency to merge, suggesting a specificity attributable to the inherent RNA sequences. This specificity in droplet behavior driven by RNA sequences could be a fundamental mechanism underlying the recognition of homologous chromosomes. Our proposed model of this is shown in Figure 6.

**Figure 6.**
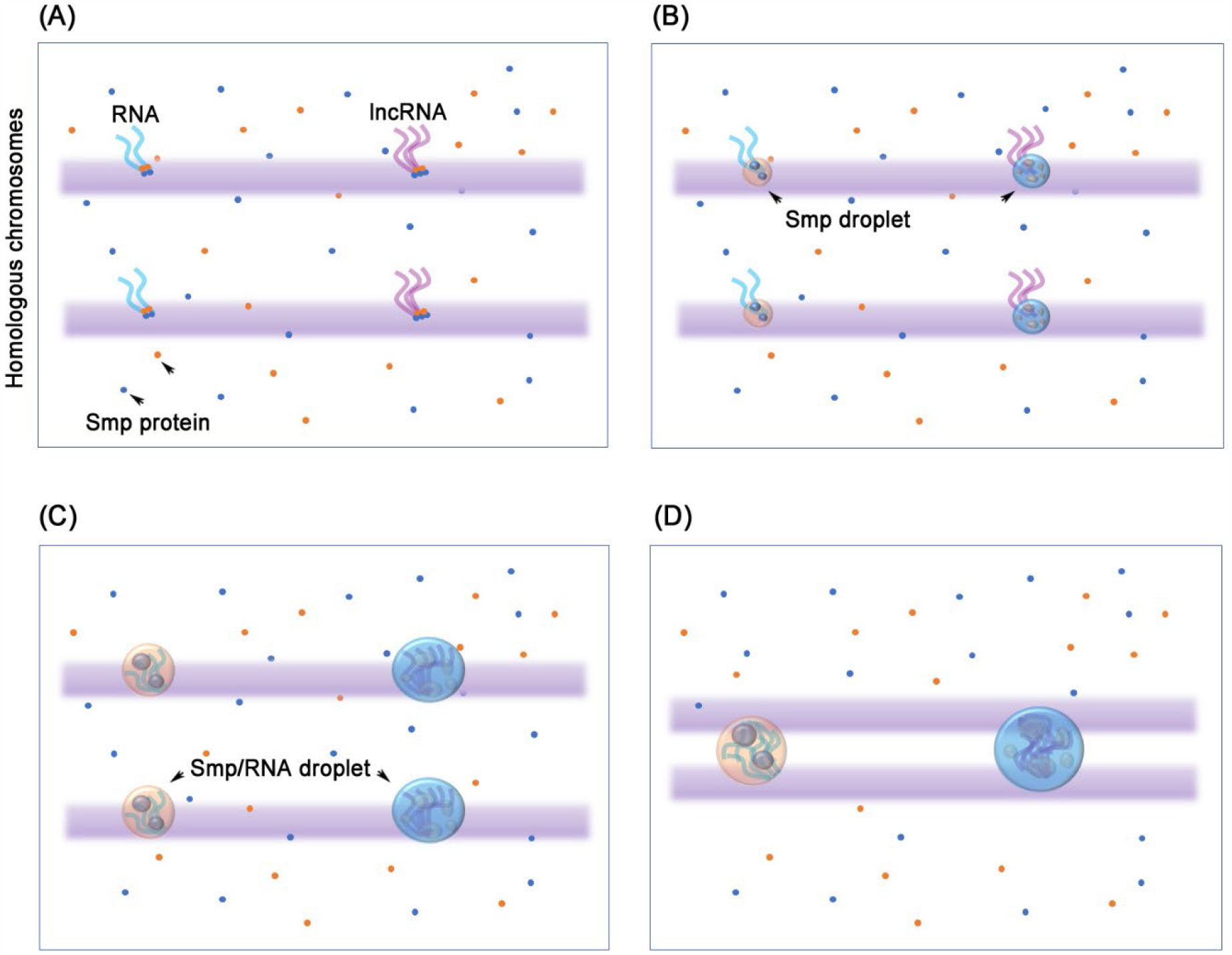
A model for Smp/RNA droplets in homologous chromosome pairing. (A) Smp proteins accumulate at the transcribed RNA and are required for transcription termination. (B) LLPS of Smps is promoted by RNA transcripts. (C) Large Smp/RNA droplets form at the transcription sites along with RNA transcript accumulation. (D) Fusions of droplets at the same allele on each homologous chromosome.

Since Smps play important role in mRNA and ncRNA transcription termination,^16^ when RNAs are transcribed from chromosomes, these Smps tend to accumulate to RNAs (Figure 6A). Elevated local concentrations of Smps and RNAs further facilitate the formation of Smp droplets (Figure 6B), the size of which increases along with the accumulation of RNA transcripts (Figure 6C). Homologous chromosomes exhibit analogous patterns of Smp/RNA droplet distribution across their surfaces (Figure 6C). When homologous chromosomes are aligned in parallel and in close spatial proximity, droplets corresponding to the same gene allele, characterized by shared intrinsic physical properties, tend to fuse (Figure 6D). This fusion process appears to facilitate the recognition and pairing of homologous chromosomes. Thus, the process of homologous chromosome pairing is likely to be achieved by amplification of the intrinsic DNA sequence into multiple RNA copies, which then congregate into Smp/RNA droplets.

In a previous study, we observed the preferential formation of Smp-RNA droplets *in vivo* at three distinct lncRNA loci,^6^ so it may be expected that coding and noncoding RNA have different roles in promoting droplet formation *in vitro*. However, with respect to the Seb1/Rhn1 droplets, we found that coding RNAs and lncRNAs promote droplet formation in a similar manner (Figure 2E). This may reflect their physicochemical properties simply in the *in vitro* assay system. In living cells, mRNA and ncRNA have different fates after transcription: transcriptional termination stabilizes mRNAs and facilitates their rapid export to the cytoplasm for protein translation. Conversely, transcriptional termination of certain categories of ncRNAs is less efficient and eventually leads to their nuclear restriction and degradation.^16-18^ In our current experiments, we used only purified RNAs and conducted *in vitro* droplet formation assays with representative Smps, which are essential for the robust pairing mechanism at the *sme2* locus. It is still unclear how the transcriptional process is involved in the droplet formation *in vivo*. Further studies will elucidate the mechanisms by which chromosome-associated RNA-protein droplets promote pairing of specific homologous chromosomes, and more generally how LLPS regulates the functional organization of chromosomes within the nucleus.

## AUTHOR CONTRIBUTIONS

Da-Qiao Ding, Jun-ichi Nakayama and Yasushi Hiraoka conceived and designed the research. Da-Qiao Ding, Kasumi Okamasa, Yuriko Yoshimura, Takaharu G. Yamamoto performed the experiments. Atsushi Matsuda set up the microscope systems. Da-Qiao Ding, Jun-ichi Nakayama and Yasushi Hiraoka wrote the manuscript with the input from all the authors.

## ACKNOWLEDGEMENT

We thank to the members of the laboratories for their assistance and discussions and Dr. Kazuhiro Aoki for providing plasmids to produce fluorescent proteins. A special acknowledgement goes to Dr. Shohei Kobayashi, whose support played a pivotal role in bringing this paper to completion.

## DATA AVAILABILITY STATEMENT

The authors declare that the data supporting the findings of this study are available within the paper and its supplementary information file.

## Abbreviations

lncRNA: long noncoding RNA;
Smp: *sme2* RNA-associated protein;
LLPS: liquid-liquid phase separation;
smFISH: single molecular RNA fluorescence in situ hybridization;
MBP: maltose binding protein;
mEGFP: monomeric enhanced green fluorescent protein;
mCherry: monomeric Cherry;
PEG: polyethylene glycol;
PMSF: phenylmethylsulfonyl fluoride;
IDR: intrinsically disordered region;
Cy3 or Cy5-CTP: Cyanine 3- or 5-labeled Cytidine triphosphate

## FIGURE LEGENDS

**Supplementary Figure 1.**
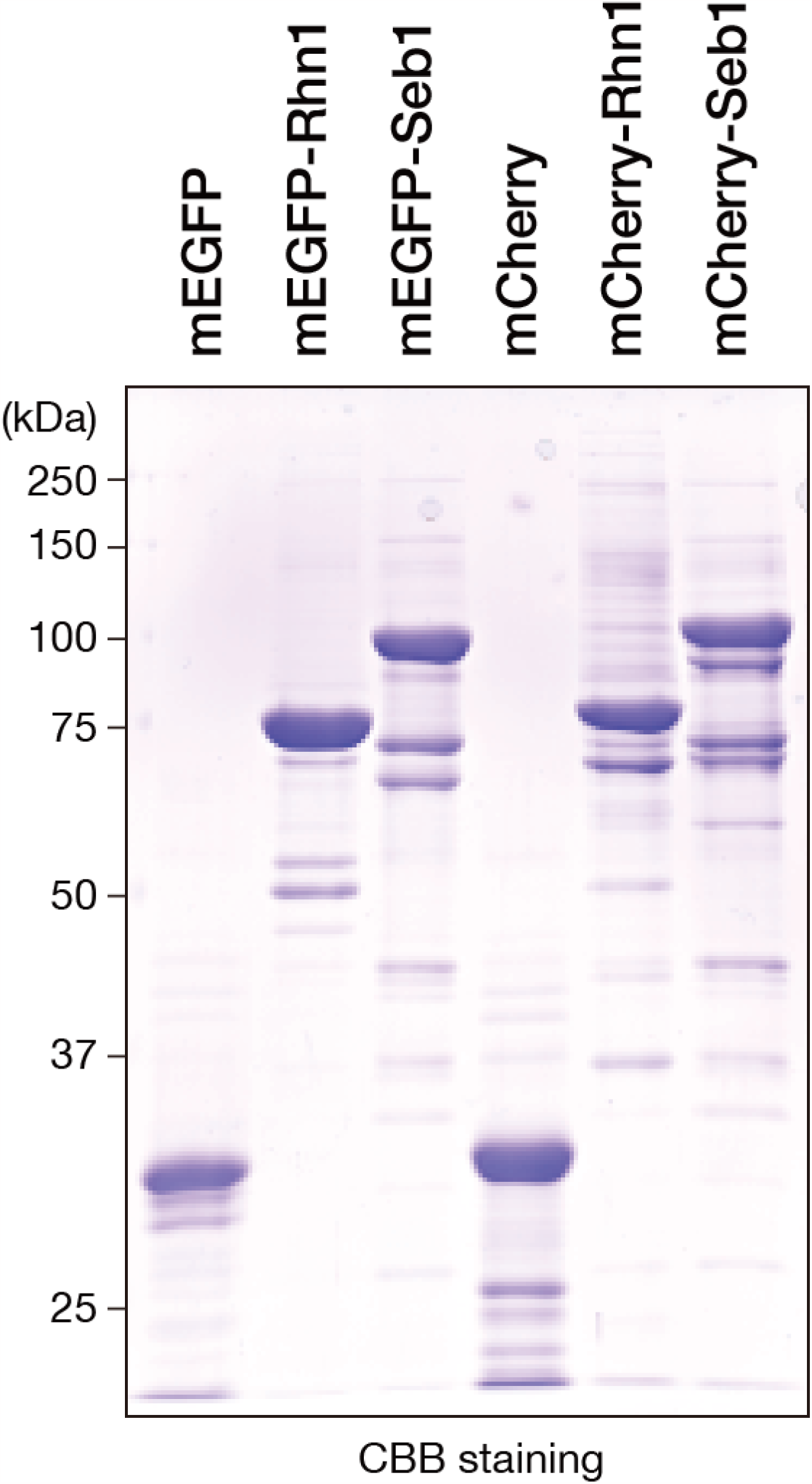
Recombinant proteins used in this study. Recombinant proteins expressed in Sf9 cells were purified, resolved by SDS-PAGE, and analyzed by CBB staining.

**Supplementary Figure 2.**
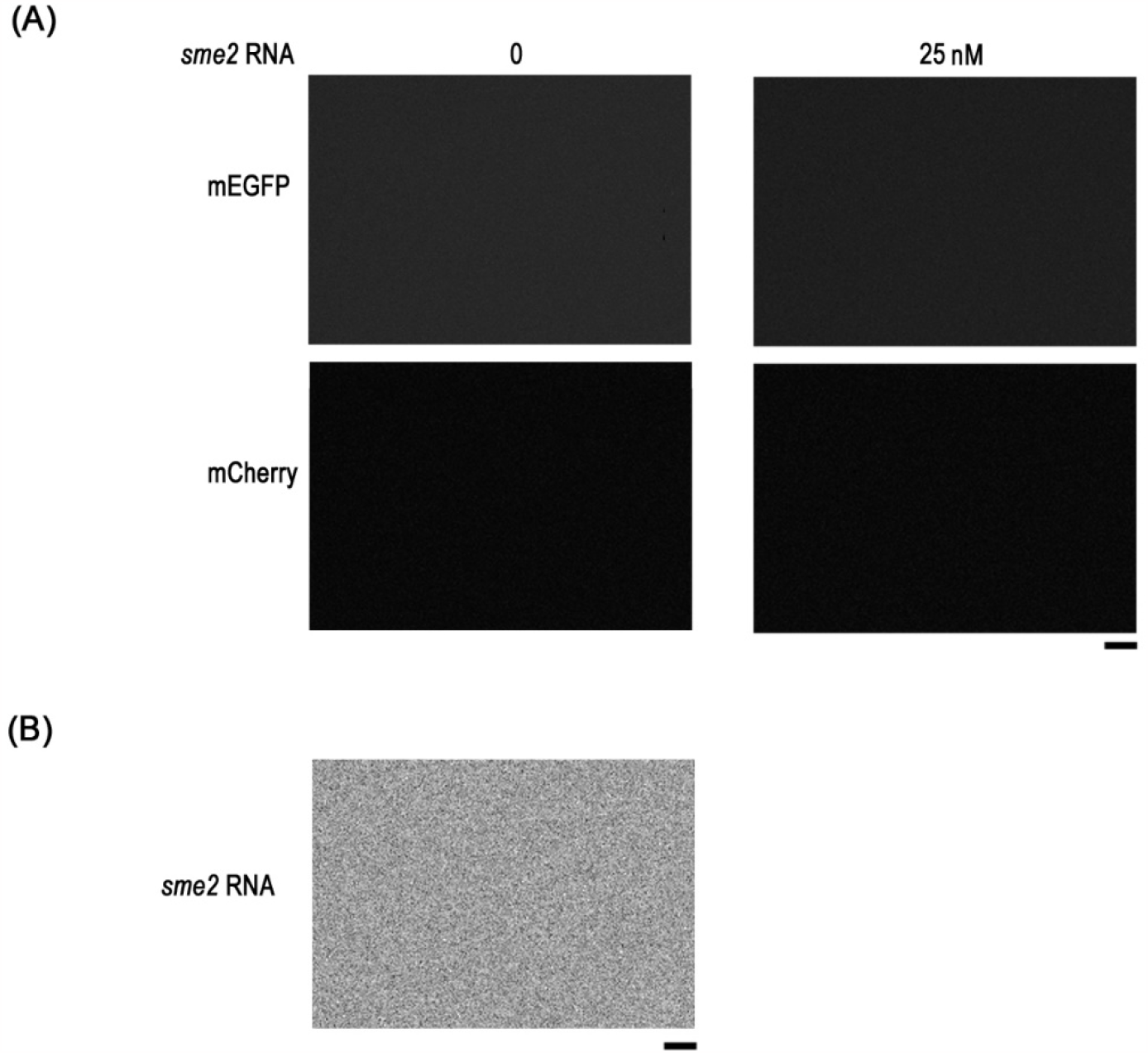
No droplet formation was found for mEGFP, mCherry, and *sme2* RNA. (A) 1 μM mEGFP or mCherry protein in phase separation buffer with or without 25 nM *sme2* RNA. (B) 25 nM *sme2* RNA in phase separation buffer. Bars, 5 μm.

